# Long-term adaptive evolution of genomically recoded *Escherichia coli*

**DOI:** 10.1101/162834

**Authors:** Timothy M. Wannier, Aditya M. Kunjapur, Daniel P. Rice, Michael J. McDonald, Michael M. Desai, George M. Church

## Abstract

Efforts are underway to construct several recoded genomes anticipated to exhibit multi-virus resistance, enhanced non-standard amino acid (NSAA) incorporation, and capability for synthetic biocontainment. Though we succeeded in pioneering the first genomically recoded organism (*Escherichia coli* strain C321.ΔA), its fitness is far lower than that of its non-recoded ancestor, particularly in defined media. This fitness deficit severely limits its utility for NSAA-linked applications requiring defined media such as live cell imaging, metabolic engineering, and industrial-scale protein production. Here, we report adaptive evolution of C321.ΔA for more than 1,000 generations in independent replicate populations grown in glucose minimal media. Evolved recoded populations significantly exceed the growth rates of both the ancestral C321.ΔA and non-recoded strains, permitting use of the recoded chassis in several new contexts. We use next-generation sequencing to identify genes mutated in multiple independent populations, and we reconstruct individual alleles in ancestral strains via multiplex automatable genome engineering (MAGE) to quantify their effects on fitness. Several selective mutations occur only in recoded evolved populations, some of which are associated with altering the translation apparatus in response to recoding, whereas others are not apparently associated with recoding, but instead correct for off-target mutations that occurred during initial genome engineering. This report demonstrates that laboratory evolution can be applied after engineering of recoded genomes to streamline fitness recovery compared to application of additional targeted engineering strategies that may introduce further unintended mutations. In doing so, we provide the most comprehensive insight to date into the physiology of the commonly used C321.ΔA strain.

**Significance Statement:** After demonstrating construction of an organism with an altered genetic code, we sought to evolve this organism for many generations to improve its fitness and learn what unique changes natural selection would bestow upon it. Although this organism initially had impaired fitness, we observed that adaptive laboratory evolution resulted in several selective mutations that corrected for insufficient translation termination and for unintended mutations that occurred when originally altering the genetic code. This work further bolsters our understanding of the pliability of the genetic code, it will help guide ongoing and future efforts seeking to recode genomes, and it results in a useful strain for non-standard amino acid incorporation in numerous contexts relevant for research and industry.

## Introduction

Billions of years of evolution has given rise to diverse organisms that share a universal genetic code. The ability to recode genomes to contain fewer than the full set of 64 triplet codons has proven useful for enabling multi-virus resistance, enhanced incorporation of non-standard amino acids (NSAAs), and capability for synthetic biocontainment. The first genomically recoded organism, *Escherichia coli* C321.ΔA (1–3), was generated by using multiplex automatable genome engineering (MAGE) (4) to substitute all 321 UAG stop codons for UAA and delete the associated Class I peptide release factor 1 (RF1), which recognizes UAA/UAG codons. Genome synthesis and assembly methods are currently being used to construct additional recoded genomes including a 57-codon *E. coli* genome (5) and a synthetic yeast genome (6–12). Given their common aim of UAG codon reassignment, these efforts would benefit from greater characterization of C321.ΔA under diverse conditions.

Previous genome engineering efforts directed toward the goal of UAG codon reassignment resulted in strains that exhibited large fitness deficits upon removal of RF1. RF1 was originally considered to be essential and only conditionally lethal mutants were described (13). In 2010, RF1 deletion was enabled by conversion of 7 essential UAG codons to UAA and introduction of a UAG suppressor (14). Two subsequent studies showed that RF1 could be deleted if Class I peptide release factor 2 (RF2), which recognizes UAA/UGA codons (15, 16), were corrected in one of two ways: By removing RF2 autoregulation (17) or by introducing a PrfB_T246A variant (18). However, the resulting strains were severely growth impaired given the presence of numerous unassigned UAG codons where ribosomes would presumably stall. More recently, RF1 could be deleted after conversion of 95 of the 273 UAG codons in *E. coli* BL21(DE3), which natively contains the PrfB_T246A variant (19); nevertheless, this strain displayed inferior growth in minimal media. C321.ΔA and its derivatives are the only strains that contain no apparent UAG codons and no RF1, enabling UAG reassignment to NSAAs without competition from off-target sites or from RF1.

C321.ΔA has experienced wide adoption as a workhorse for NSAA incorporation. NSAAs broaden the repertoire of biological chemistry in living systems for diverse purposes such as photocrosslinking (20, 21), functionalization (22), structure determination (23), fluorescence (24, 25), metal binding (26), biosensing (27), and immobilization (28). NSAAs also augment protein function, such as the affinity and pharmacodynamic properties of therapeutically relevant proteins (29–31) and the catalytic properties of industrially relevant enzymes (32–34). Many of these potential applications for NSAAs, such as live cellular imaging, metabolic engineering using simple carbon sources, and industrial-scale protein expression, necessitate the ability to culture cells in defined media. However, C321.ΔA exhibits a fitness deficit compared to its non-recoded ancestor, which is due at least in part to off-target hitchhiker mutations that accumulated during recoding (4). This fitness deficit is exacerbated in defined media.

Experimental evolution is a powerful method for directly observing rapid evolutionary change in the laboratory, and non-model organisms can quickly adapt to lab conditions after a period of sustained propagation (35). We describe here the first example of ALE of an organism containing a genome with fewer than 64 codons. By sequencing the whole genomes of two clonal isolates from more than 50 independent evolved populations, we present evidence that the conversion of 321 UAG codons into UAA codons, and especially the subsequent deletion of RF1, introduces a burden to *E. coli* K-12 cellular translation machinery under a wide range of industrially relevant defined media. Our evolutionary analysis reveals that point mutations in RF2 that are known to provide increased activity on UAA codons are selected for and recover much of the fitness loss. Furthermore, we observe that natural selection exhibits a variety of mechanisms to correct the most detrimental off-target mutations introduced during engineering of the recoded strain, including the use of premature termination codons (PTCs) in essential genes and the inactivation of a stress related transcription factor.

## Results and Discussion

### Adaptive evolution achieves robust growth of evolved recoded strains in glucose minimal media

We began by seeding 14 independent populations for each of the following four strains: (i) ECNR2 (the non-recoded parent strain of C321.ΔA); (ii) C321.ΔA (a C321 derivative with RF1 removed); (iii) C321.ΔA-v2 (a C321.ΔA derivative containing engineered reversions to three off-target MAGE mutations) (36); (iv) C321 (a C321.ΔA derivative with RF1 restored to its native locus). These strains contained inactivated *mutS*, which significantly increases MAGE efficiency but also results in a hypermutator phenotype. Hypermutators are known to arise naturally during long-term evolution (37), and we hypothesized that we might accelerate our ALE by using MutS-strains for seeding. Independent lineages were propagated for over 1,000 generations and subject to genotyping, doubling time analysis, and storage at approximately 70-100 generation intervals (**Fig. 1**). We increased the dilution and passaging rate as growth rates increased so that strains did not evolve to survive long periods of stationary phase, but instead experienced selective pressure primarily for exponential growth. The decision to passage in batch culture rather than using continuous cultures is one with trade-offs (38), but was made both in light of our desire to replicate normal laboratory conditions, which will include periods of stationary phase, and the more practical considerations involved with building more than fifty chemostats.

**Figure 1.**
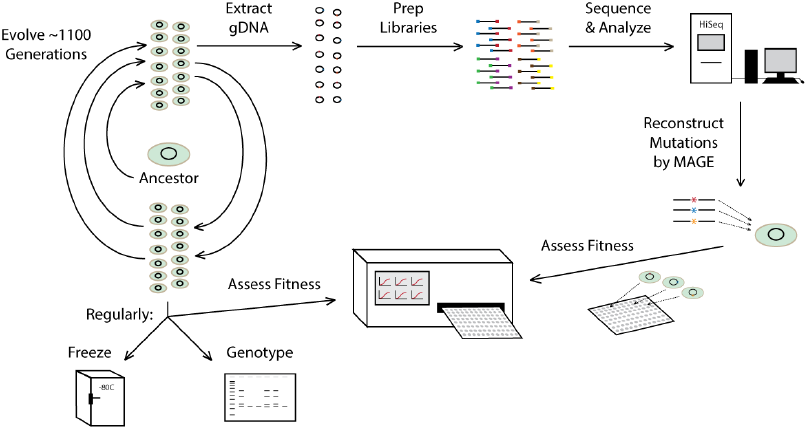
Project workflow for long-term adaptive laboratory evolution (ALE) of recoded *Escherichia coli*. Briefly, strains were passaged in 14 independent replicates for 1,000+ generations with routine genotyping, storage, and growth rate analysis by plate reader. Two clones from each population at the final time point were used for genomic DNA preparation and Illumina sequencing library preparation. Sequencing was performed, data was analyzed to identify variants, and MAGE was used to reconstruct alleles in ancestralstrains.

After ~1,100 generations we determined, through intermittent sampling of population doubling times, that the fastest populations from both the experimental and the control lines were approaching, in minimal media, the doubling time of ECNR2 in rich media (**Fig. 2**). We therefore paused the evolution at this point and characterized and sequenced all evolved populations. While the recoded strains experienced the largest changes in fitness, all experimental populations converged upon a similar doubling time at around 40 minutes for the fastest growing biological isolates. Coarse samplings of the doubling time improvements over evolutionary time was run on all evolved populations (**SI Fig 1**). We then chose two populations from each strain, except for C321.ΔA-v2 for which we chose four populations, which showed the greatest improvement to doubling time by the end of the evolution, and measured the growth rate improvement over time more finely (**Fig. 2**). All strains showed marked improvement to their fitness in minimal media over the time course of the ALE. Of note is that the C321.ΔA-v2 lines maintained a small fitness edge over the C321.ΔA lines throughout the experimental course, although the C321.ΔA lines did narrow the gap. On the other hand, the C321.ΔA lines caught up in fitness to the C321 lines, which had had RF1 restored to its native locus, which could imply that the fitness cost of RF1 deletion from C321 was readily mitigated by ALE. Next, we sought to identify any causal mutations behind these improvements to fitness from an analysis of whole genome sequencing of two clonal isolates from each evolved population.

**Figure 2.**
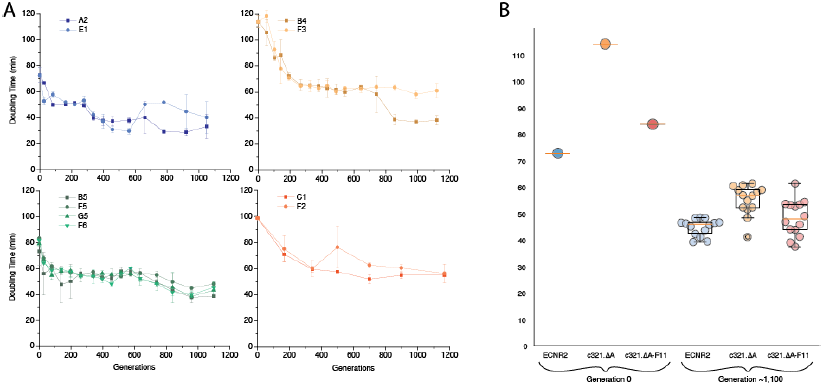
Representative fitness trajectories during evolution for different ancestralstrains. (A) Top-Left: Two lineages of non-recoded ECNR2 (engineered from *E. coli* MG1655 K-12). Top-Right: Two lineages of recoded C321.ΔA (321 UAG->UAA & RF1-). Bottom-Left: Four lineages of recoded C321.ΔA-v2 (C321.ΔA with engineered reversion of some off-target mutations that occurred during recoding). Bottom-Right: Two lineages of recoded C321 (C321.ΔA with *prfA* gene restored). (B) Ancestraland final doubling time measurements sampled from all lineages of ECNR2 (blue), C321.ΔA (yellow), and C321.ΔA-v2 (green).

### Stop codon identity in evolved strains displays differential selection by population phenotype

As recoded C321 has a globally-altered stop codon repertoire, we first chose to examine how ALE differentially affected stop codons in the evolved populations. We hypothesized that RF1 removal from C321 may put selective pressure on stop codons during adaptive evolution. The effectiveness of stop codons at coding for translation termination is affected by their identity (UAG, UGA, or UAA) and by sequence context (39–41). Suppression experiments suggest that UAA codons, which are recognized by both RF1 and RF2, are read more often by RF1 in *E. coli* K12 (42). Analyses in *E. coli* show that codon adaption index correlates positively with increasing UAA preference (43) and that most highly expressed genes end in UAA (44). In the absence of RF1, insufficient RF2 activity in C321.ΔA strains may impair fitness due to ribosome stalling at UAA or UGA stop codons. This would result in a smaller pool of free ribosomes, lower translational capacity, and increase the requirement for ribosome rescue by tmRNA-mediated trans-translation (45, 46) or by alternative ribosome-rescue factor (47, 48).

We examined all stop codons in sequenced clones to determine whether they changed identity. We considered three potential events: (i) direct conversion from one stop codon to another; (ii) read-through effects of mutating a stop codon into a sense codon; and, (iii) frameshift mutations close to the end of a protein-coding gene which would have the effect of switching termination from the wild-type stop codon to a new position. We only considered frameshift mutations that both occurred within 100bp (~33 amino acids (AA)) of the 3’ end of a gene and that did not add more than 200bp (~66 AA) of additional length to the end of a gene.

We saw only three instances of direct UAA to UAG reversions, indicating a robustness of the recoding performed to remove UAG stop codons from the strain to long-term laboratory passaging. The three cases of UAA to UAG reversions that we did observe were to *hemA* in C321 during the re-introduction of *prfA* to the strain in preparation for long-term passaging, and two reversions to the pseudogenes *eaeH* and *ychE* during passaging in two clones from a C321 (*prfA*+) lineage and one clone from a C321.ΔA-v2 lineage. We exclude these two latter reversions from our downstream analysis of changes to termination codons because they both occur in pseudogenes, and so are unlikely to be of great significance. Using the broader criteria described above, we see 39 unique instances of stop codon changes in this work (**SI Table 1**). Three changes are excluded from further analysis because they occur in pseudogenes, and five changes to *rph* and one to *wzzE* are ignored because they are either direct or indirect repairs of prior frameshift mutations occurring in either the wild-type K-12 strain, in the case of *rph*, or the C321 ancestor during *prfA re-introduction*, in the case of *wzzE*, This leaves 30 total termination codon changes, 9 of which are due to a point mutation to the termination codon itself, and 21 of which are due to frameshift mutations (**Fig. 3**).

**Figure 3.**
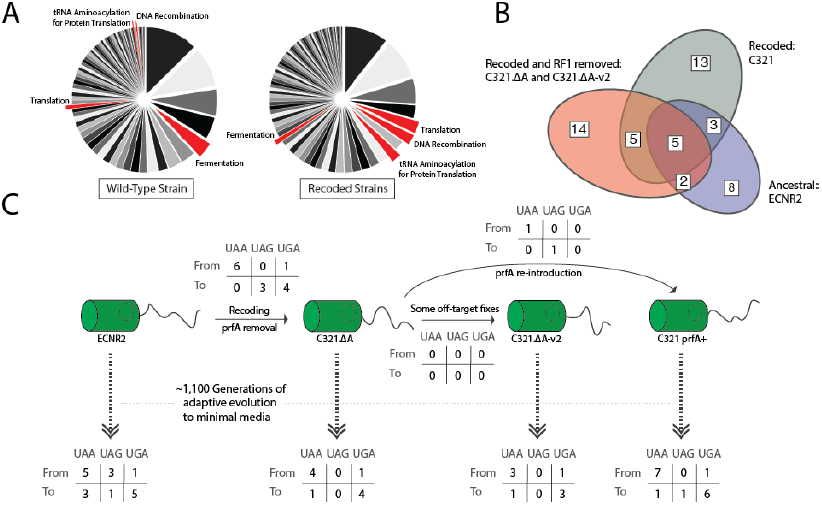
Next-generation sequencing results and variants of interest from independent evolved populations. (A) Gene ontology results showing all categories of mutations found in non-recoded ECNR2 (left) or in recoded strains (right). Highlighted in red are categories of interest. (B) Venn diagram showing distribution and quantity of mutated genes found in individual strains or shared among strains. (C) Results from analysis of changes in stop codon usage in evolved lineages.

Two trends stood out to us in inspecting the data from the results of the sequencing of LTE populations. First, there is a general trend that in all populations, including those with RF1 present, genes with UAA codons experience more mutation than genes with UGA codons. Wild-type MG1655, in the latest annotated genome from the NCBI, has UAA:UGA:UAG codon ratios of approximately 10 to 5 to 1 respectively. In this study, however, we see that genes with UAA codons see about four to five times as many termination codon alterations as genes with UGA codons (about twice the expected rate P < 0.1) and that genes with UAG codons are also disproportionately targeted for termination codon alteration during ECNR2 adaptive evolution (all other populations are C321 derivatives with UAG codons removed). Furthermore, most of the stop codon reassignment seen is due to either frameshift mutations near the 3’ end of protein-coding genes or read-through of termination codons by way of their mutation to sense codons (**Table S3**). Theoretically these two causes should show an equal propensity to mutate to any of the three termination codons, but we observe a significant bias toward UGA stop codons. These observations seem to indicate an adaptive preference for UGA over UAA or UAG in minimal media.

The second notable trend in termination codon reassignment is that at every transition between ECNR2 and the various C321 derivative strains there was at least one mutation toward a UAG codon except in those cases where RF1 was absent. This seems to be a clear indication of the fitness cost of mutation toward a termination codon in a strain missing its cognate release factor. Lastly, notable perhaps for its absence, is that no increased mutational load on stop codons overall is noticed in recoded lineages as compared to the wild-type ECNR2 lineage, even in the absence of RF1. This suggests that stop codon reassignment is not an easily accessible source for fitness gains to the recoded C321 strains.

### Selective mutations occur in translational machinery: *prfB* and *prfC*

Three mutations observed repeatedly were found in the *prfB* and *prfC* genes. *PrfB* encodes RF2, and *prfC* encodes release factor 3 (RF3), which is a ribosome-dependent GTPase that stimulates release of RF1 and RF2 from the ribosome after termination (49–51). In the recoded lineages missing *prfA* (C321.ΔA and C321.ΔA-v2), we observe independently-arising missense mutations to the remaining release factor: *prfB*. The dominant mutation, observed in 31 out of 54 sequenced clones from *prfA*-lineages, is PrfB_T246A, which is a revertant to the PrfB sequence present in most non-K-12 *E. coli* strains (52). This mutation has been characterized *in vitro* (52) and *in vivo* (53, 54), and it is known to increase net RF2 activity on UAA codons by roughly 5-fold *in vivo* (54). Another mutation observed in independent lineages is PrfB_E170K. Based on previously targeted mutagenesis studies of *prfB*, PrfB_E170K appears to also have increased termination efficiency for UAA codons (55, 56). We also observe five separate missense mutations to *prfC* in the recoded lineages, but in contrast to *prfB*, for which mutations only appear in the recoded lines missing *prfA, prfC* mutations appear also in the recoded C321 line with *prfA* reintroduced. The most frequently-observed mutation to *prfC* is PrfC_A350V, which arises independently in at least one population from all three recoded lines. To our knowledge, no one has studied the effect of the A350V mutation, although this position was one of many targeted for mutagenesis in a previous study (57). During a study of strains containing temperature sensitive RF1 and RF2 variants, *prfC* mutations appeared at positions 96, 118, 399, and 440, each of which suppressed growth defects (58). The previous observation of RF3 mutations during defective translation termination lends support to the notion that the A350V mutation may also alleviate impaired termination. Note that when translation termination is impaired in other ways, such as by deletion of ribosomal modification machinery, mutations in *prfB* and in *prfC* are also known to arise, but not at the positions observed in our study (see Supplemental Discussion). Overall, in our study, three RF mutations were found to occur in independent lineages: PrfB_E170K, PrfB_T246A, and PrfC_A350V. None of these mutations were found in evolved ECNR2 populations. Furthermore, no sequenced clones were found to contain more than one of these three mutations, and a functional RF1 seems to be enough to epistatically shield *prfB* from selectional mutation.

### The contribution of mutated RF alleles to fitness depends on media composition

We reconstructed these RF mutations individually in ancestral strains and tested their effects on fitness using two approaches: doubling time analysis and head-to-head competition. We also examined the effect of each mutation in a wide range of relevant media conditions. Media composition has been shown to influence how impaired translation termination affects growth rate (See Supplemental Discussion). In general, translational termination defects often show increased severity in poorer media conditions due potentially to two factors: (i) slower-growing *E. coli* cells produce lower levels of release factors and fewer ribosomes, meaning more demand for release and recycling on fewer molecules (59), and (ii) growth in minimal media or on low quality carbon sources necessitates expression of a broader set of genes (60, 61), which more often contain weak termination codons than highly expressed genes (43). We therefore decided to ask whether adaptive fitness improvements to RFs would be applicable across media types.

Although we performed LTE only in M9+glucose, we measured doubling times for strains grown in three defined media (M9+glycerol, M9+glucose, and MOPS EZ Rich) and three complex media (LB, LB+glucose, and 2XYT) (**Fig. 4**). To complement the doubling-time analysis, we performed head-to-head competition assays against a fluorescent reference strain (**SI Fig. 2)**. Several trends emerged from this data. First, in agreement with past observation, we saw that doubling times were much larger for ancestral recoded strains than for ECNR2 across media types. Next, in support of the idea that RF stress is more acute in poorer media conditions, we observe that allelic fitness variants have little effect in rich media, but offer large improvements to recoded strain fitness in defined media. Furthermore, in accordance with RF variants being the most acutely selected variant type in this study, we observe a decreasing difference in fitness between recoded and non-recoded strains as media becomes richer, implying that RF stress is a key driver of fitness loss in minimal media for the recoded strains. Overall some RF mutations were beneficial in defined media but neutral in complex media; the mutation that consistently appeared most beneficial to recoded strains was PrfB_T246A, which is corroborated by its greater frequency of occurrence during evolution.

**Figure 4.**
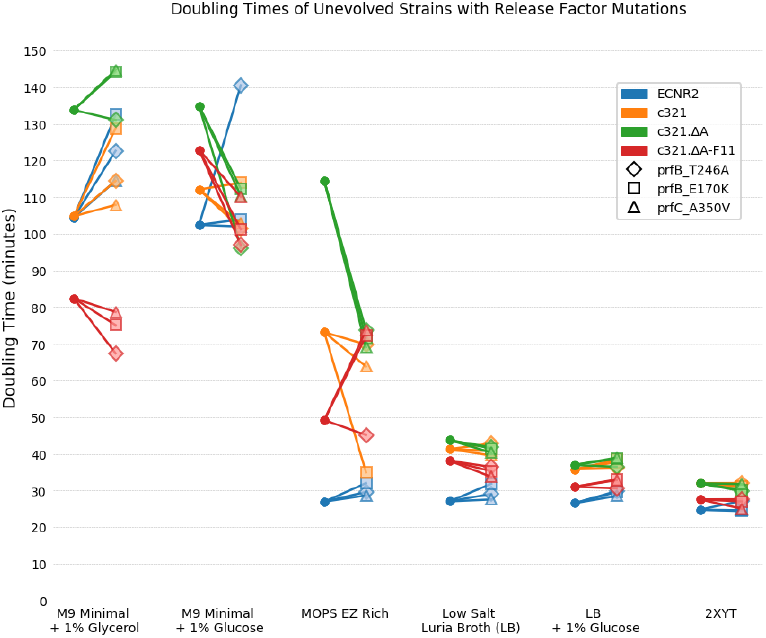
Effects of individually reconstructed release factor mutations and media context on growth rate of ancestral strains. Different media compositions are shown on the x-axis arranged from poorer carbon sources in defined media (left) to rich and complex media (right). The three release factor mutations investigated are PrfB_T246A, PrfB_E170K, and PrfC_A350V.

### Selective mutations occur in genes not associated with translational machinery

In addition to the RF mutations described above, we conducted a genome-wide analysis looking for genes mutated multiple times across independent lineages. As a threshold for significance, we chose to set the bar at genes for which we see at least five mutant alleles in one of the four parental lineages. We chose this threshold to ensure that at least three separate instances of mutation to the gene were recorded, as two clonal isolates were sequenced form each evolved population. There were 52 genes that cleared this bar (**Table S2**), many of which we were able to identify as being commonly hit genetic targets from previous ALE studies or as being otherwise characterized in the literature. In particular, many of these 52 genes were found previously during adaptive evolution of *E. coli* K-12 strains to minimal media. These include pykF, *rph, rpoB*, and *rpoC*, found in three studies from the Palsson laboratory (62), and *fis, gltB, kup*, and again *pyk F* found by the Lenski laboratory (63). Many of these gene targets have been discussed, but our study does provide some further insight in several cases.

It has been previously documented that a frameshift mutation in the C-terminus of *rph* in wild-type K-12 strains causes pyrimidine starvation in minimal media. This is due to low *pyrE* expression because a premature *rph* stop codon impedes transcription/translation coupling, which supports optimal levels of *pyrE* expression past the intercistronic *pyrE* attenuator (64). Evolutionary studies from the Palsson lab identified a recurring 82-bp deletion in the regulatory region between *rph* and *pyrE*. In contrast, we in no cases see this same deletion, but instead see thirteen separate mutations to a 233-bp stretch extending from the C-terminus of *rph* through some intergenic space, and into the first 15-bp of the *pyrE* gene. The five most frequent mutations make up 86% of the total hits. Four of these target the first attenuator stem, weakening it (**SI Fig. 3**), and the other, found in eight clonal isolates from four populations, is an exact reversion (3815879 TCC to TCCC) to restore the original wild-type locus found in the K-12 ancestor. Other mutations found are frameshifts in *rph* to extend its reading frame either by 10 amino acids to its wild-type UGA or a further 14 amino acids to a UAG stop codon, which we presume also relieves *pyrE* attenuation.

Of the other genes that we found in common with previous ALE studies, mutations to RNA polymerase subunits B and C (*rpoB* and *rpoC*) have been particularly well-documented. In one study, Conrad et al. characterized three *rpoB* mutations that were found during ALE to glucose minimal media, and found that all variants were adaptive in defined media, but were less fit in rich media (62). Conrad et al. additionally found that these variants exhibited decreased open complex longevity and greater processivity, which they proposed led to greater expression of metabolic genes, and lower rRNA expression. In this work we found eleven unique *rpoB* variants and 20 unique *rpoC* variants, all of which are mis-sense mutations, and none of which were documented in previous ALE studies. This suggests to us that the landscape for adaptive fitness gain in the two polymerase subunits is quite large, with many routes to switching from LB adaptation to M9 adaptation.

Of the remaining genes unrelated to translation that were selectively mutated in evolved recoded populations, we chose five of those most over-represented in recoded strains for allelic reconstruction in the C321.ΔA-v2 and C321.ΔA-v2.PrfB_T246A backgrounds. We chose to assay in these backgrounds to ask whether there were additive fitness effects from non-translational mutations. With one exception (*fimH*), these genes all contained premature termination codons (PTCs) or frameshifts that appeared in different regions of the gene. Therefore, in these cases we introduced PTCs approximately 30 bases downstream of the start codon during allele reconstruction, whereas in *fimH*, we tested the two most frequent missense mutations. When we compared the fitness of the reconstituted variants by competition assay with the unaltered strains, C321.ΔA-v2 was improved by every mutation, whereas the fitness effects in C321.ΔA-v2.PrfB_T246A were muted, with only FimH_S83L providing a significant fitness benefit (**SI Fig. 4**).

In only our evolved recoded strains, we observed eighteen mutations, including PTCs appearing in the *folA* gene, many of which occur in the promoter region which has an off-target hitchhiker mutation from the recoding of C321 (C49765T). The four most frequent mutations appear in the promoter region near the hitchhiker mutation. We also observe four separately-arising PTCs, and many low-frequency mutations later in the gene or to the 5’ end of *folA*, in a region likely to impact its mRNA structure (65). Interestingly, after alternating the passaging of C321.ΔA in LB and minimal media for just 12 growth cycles and sequencing five clones, Monk et al. similarly observed one instance of a PTC in *folA* and two mutations in the promoter region (66). Given the promoter mutations, Monk et al. predicted that these mutations lead to an increase in metabolic flux through this enzyme. However, we viewed the PTC mutations in *folA*, an essential gene, as a likely signature for downregulation of FolA protein levels, which we thought would be favored if the hitchhiker mutation resulted in higher than wild-type *folA* expression. Indeed, nonsense mutations in *E. coli* genes have been known to not completely abolish corresponding enzyme activity, instead allowing residual expression of roughly 10^−4^ of wild-type levels (67–71).

To investigate this hypothesis, we cloned the native and C321.ΔA *folA* promoter sequences upstream of a fluorescent reporter construct and measured fluorescence over a 24-hour time course in M9+glucose and LB media (**Fig. 5A,** and **SI Fig. 5**). We also measured fluorescence in control strains containing no reporter construct or a moderately strong synthetic constitutive promoter. Strains expressing fluorescent reporter using either the C321.ΔA *folA* promoter or the strong constitutive promoter exhibited a steadily increasing and high level of fluorescence in both media. In fact, fluorescence resulting from the C321.ΔA *folA* promoter exceeded fluorescence resulting from the constitutive promoter for much of the time course. On the other hand, fluorescence was essentially undetectable for the native *folA* promoter (same as the no reporter control) in both media. This result provides strong evidence in support of the hitchhiker mutation in the *folA* promoter significantly increasing FolA expression. Given that fluorescence signal from the native *folA* promoter was so low under both media conditions, it is likely that the *folA* expression in C321.ΔA is globally burdensome and it should be corrected through engineering or evolution for all envisioned applications.

**Figure 5.**
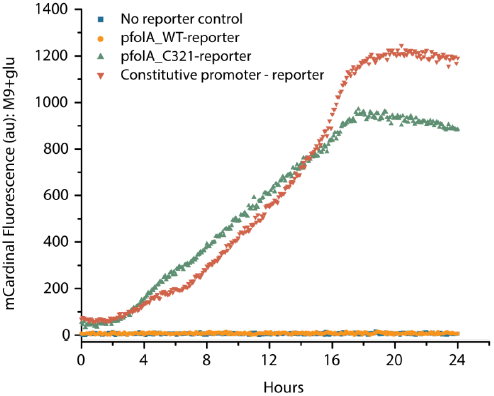
Effect of the *folA* promoter mutation on fluorescence resulting from fluorescent protein gene expression.

### Evolved recoded strains are capable of NSAA incorporation and most useful for defined media context

The absence of UAG codons and RF1 permits dedicated reassignment of UAG to an NSAA. To evaluate the utility of our evolved recoded strains for NSAA incorporation, we isolated three colonies and co-transformed them with plasmids harboring an orthogonal translation system (OTS) and a reporter protein containing UAG sites for NSAA incorporation. Specifically, we used the bipyridylalanine aminoacyl-tRNA synthetase (BipyARS) and *Methanococcus jannaschii* tRNAopt_CUA_ as our OTS, which was recently shown to be more orthogonal to the native 20 amino acids than related engineered *M. jannaschii* tyrosyl-tRNA synthetase variants (72). We chose two fusion proteins of ubiquitin and super-folder GFP (Ub-sfGFP), one containing no UAG sites as a positive control for protein expression and the other containing 2 UAGs at positions between the two domains and internal to the GFP (72). We measured GFP fluorescence normalized to culture optical density (FL/OD signal) in the presence and absence of p-Acetyl-phenylalanine (pAcF), which is an inexpensive NSAA known to be a substrate of BipyARS (**Fig. 5A**). To avoid catabolite repression of the arabinose inducible promoter system that regulates BipyARS, we used glycerol minimal media for these experiments.

Upon testing expression of the 0UAG reporter, we observed FL/OD signal unaffected by the addition of pAcF for ECNR2 and a representative evolved recoded strain. The C321.ΔA strain harboring both plasmids did not observably grow in glycerol minimal media during a 24-hour observation period, even without induction for either plasmid-based promoter system. Notably, the FL/OD signal corresponding to the evolved recoded strain was ~20-fold less than that of ECNR2. When we tested NSAA incorporation using the 2UAG reporter, we observed substantially lower FL/OD signal consistent with the competition for UAG suppression with RF1 in ECNR2 and the limited activity of previously developed orthogonal translation systems even in the absence of RF1. Interestingly, the FL/OD signal in the presence of pAcF did not vary significantly between cultures containing ECNR2 or evolved recoded strains, but the undesired signal in the absence of pAcF was lower for cultures containing evolved recoded strains compared to ECNR2. This result demonstrated that recoded strains can not only grow and incorporate NSAAs in minimal media after evolution, but they can also exhibit better dynamic range for NSAA incorporation applications.

Our results demonstrate that the capacity for target protein overproduction in glycerol minimal media, at least for GFP using a pZE21 vector, has substantially declined for evolved recoded strains compared to the non-recoded ancestor ECNR2. Based on our data, recoded strain capacity for protein overproduction in this medium primarily improved due to fitness gains resulting from evolution. However, our evolution scheme would not have favored protein overproducers, and rational genome engineering can be used to further improve protein expression, potentially even at the cost of some fitness. Well-documented targets include proteases encoded by *lon* (73) and *ompT* (74), as well as nucleases encoded by *rnb* (75), *rne* (76), and *endA* (77). Others have attempted to engineer some of these targets in C321.ΔA and observed increased protein expression (78). While the expression of NSAA-containing proteins may also benefit from these approaches, NSAA incorporation faces other limitations that ought to be addressed, such as limited activity and selectivity in the engineered orthogonal aminoacyl-tRNA synthetases used to introduce NSAAs into proteins. An unusually selective but not highly active synthetase was used for NSAA incorporation in this study. Rational engineering or directed evolution approaches can be used to generate synthetases with improved attributes in either or both categories.

Finally, we measured the doubling times of a previously engineered strain, C321.ΔA.fix, with the C321.ΔA.M9adapted strain developed through ALE in this study. Not surprisingly, C321.ΔA.M9adapted exhibited much faster growth than C321.ΔA.fix in glucose minimal media (**Fig. 6C**). Our result shows that while genome engineering strategies and laboratory evolution are both useful tools, they each present tradeoffs that suggest different use cases and order of operations. Genome engineering strategies facilitate the making of rational changes once targets or designs are clearly identified or known. However, these approaches can also introduce undesired changes (1), and they may also assume that identified designs are globally optimal. In contrast, laboratory evolution enables natural selection to guide both steps of target identification and change-making at once, and it is the most logical tool for improving the phenotype of fitness. It is particularly beneficial when applied after large-scale genome engineering to evaluate the robustness of engineered alterations and to correct unwanted alterations. However, evolution performed in specific contexts can result in specialized strains that are less fit in other contexts (62). Evolution is also limited in the total number of traits, and the simultaneous number of traits, that it can be used to enhance because of the selection requirement (79). Thus, these strategies should be used in concert.

**Figure 6.**
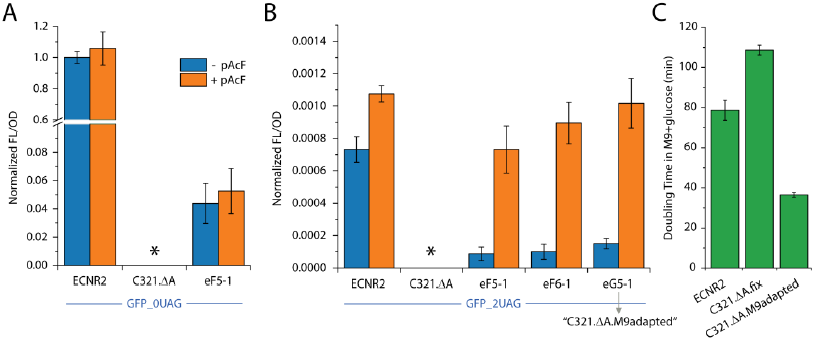
Characterization of select evolved recoded clones for applications featuring NSAAs. (A) Expression of a control GFP reporter containing no UAG codons in various strains in the presence and absence of the NSAA p-acetyl-phenylalanine (pAcF). (B) NSAA incorporation assay based on expression of a GFP reporter containing 2 UAGs in various strains in the presence and absence of pAcF. Evolved recoded clone G5-1 was selected as “C321.ΔA.M9adapted” for deposit in the Addgene repository. (C) Doubling time measurements for C321.ΔA.M9adapted and related strains in glucose minimal media.

## Conclusions

Here we overcome one of the principal shortcomings of the recoded C321 lineage, evolving a derivative strain that grows robustly in both rich and minimal media, and show that adaptive evolution is a useful tool for the recovery of fitness loss from laboratory engineering. After only ~1,100 generations of adaptation to minimal media, independent C321 lineages recovered nearly all of their fitness loss from the recoding of 321 stop codons and the correlated off-target mutational load. Previous efforts with adaptive evolution in *E. coli* K-12 strains that targeted growth in minimal media have resulted in superior growth rates in minimal media but inferior growth rates in rich media compared to the parent strain (62). In this work, we observe superior growth for evolved recoded strains in all defined media and no significant difference in rich media. The hypermutator phenotype may have sped the evolutionary trajectories of these strains, but without apparent ill-effect on the final cell lines. The evolved lines still show somewhat diminished protein expression, but show great improvement over the original recoded strain, and have a cleaner NSAA incorporation profile than wild-type ECNR2. We anticipate that the deposited strain C321.ΔA.M9adapted will be of significant interest to researchers interested in a fast-growing C321 derivative that thrives in a broad range of media types. Particularly compelling use cases for C321.ΔA.M9adapted include recombinant protein expression in defined media, cellular microscopy, and metabolic engineering.

## Materials and Methods

### Strains

The following four strains were each passaged in 14 independent populations for approximately 1100 generations in defined M9 minimal media + 1% glucose + biotin + carbenicillin: ECNR2 (4), C321.ΔA (1), C321.ΔA-v2 (36), and C321 (*prfA*+). C321.ΔA-v2 is a variant of C321.ΔA in which some off-target mutations were corrected using MAGE. C321 was constructed by scarlessly reintroducing the *prfA* gene into its wild-type locus in C321.ΔA using recombineering and CRISPR-Cas9-based selection (80). In brief, the *prfA* gene along with flanking homology was amplified from ECNR2 to generate a linear DNA cassette for recombination. C321.ΔA was transformed with plasmids containing inducible Cas9 and guide RNA containing the junction sequence at the site of insertion. C321.ΔA was heat shocked to induce the lambda red system as in standard recombineering/MAGE protocols and transformed with the *prfA* cassette for homologous recombination. All of these strains are derived from *E. coli* K-12 MG1655.

### Culture Conditions

A 1X M9 salt medium (M9+glucose) containing 6.78 g/L Na_2_HPO_4_ 7H_2_O, 3 g/L KH_2_PO_4_, 1 g/L NH_4_CI, and 0.5 g/L NaCl, supplemented with 1 mM MgSO4, 0.1 mM CaCl_2_, 1% glucose, trace elements, biotin, and carbenicillin was used as the culture medium for evolution and most experiments in this study. The trace element solution (100X) used contained 5 g/L EDTA, 0.83 g/L FeCl_3_·6H_2_O, 84 mg/L ZnCl_2_, 10 mg/L CoCl2·6H_2_O, 13 mg/L CuCI_2_·2H_2_O, 1.6 mg/L MnCl_2_·2H_2_O and 10 mg/L H_3_BO_3_ dissolved in water (81, 82). Trace element solution was added to a concentration of 1X to decrease variability due to metal content in water.

The following media are also used in this study: (i) M9+glycerol medium, which is identical to M9+glucose except with 1% glycerol substituted for 1% glucose; (ii) MOPS EZ Rich medium, which is a commercially available defined medium (Teknova, Cat. No. M2105); (iii) Low Salt (0.5g/l NaCl) LB Lennox (LB) medium (; (iv) Low Salt LB supplemented with 1% glucose (LB + glucose). Adaptive evolution

For adaptive evolution, ancestral strains were streaked on LB agar plates, and colonies were used to inoculate 14 independent 1 mL culture volumes in glucose minimal media. Cells were typically passaged during late exponential phase at a dilution of 1:1,000 in 1 mL culture volumes incubated in deep 96-well plates at 34 C and 400 rpm. The dilution and passaging rate varied with the strains’ growth, meaning that strains were initially diluted between 1:100 and 1:500 once per day (~7-9 generations per day), and then gradually ramped up to a maximum passaging rate of two 1:1,000 dilutions each day (~20 [log2(2,000)] generations per day as their fitness increased.

### Sequencing library preparation

After evolution, two clones were isolated from each population for genomic DNA purification and for next-generation sequencing using an adapted form of the Illumina transposase library preparation (83). Barcoded libraries were pooled and run in an Illumina HiSeq 2000.

### Genome resequencing and analysis

Genome resequencing and analysis was performed in two different ways to focus on either evolved variants of gene sequences or on individual single nucleotide polymorphisms (SNPs) that emerged: (i) using Millstone (84); (ii) using a custom analytics pipeline. (i) Millstone is a custom software suite built by our lab for the purpose of whole genome mutational analysis. Illumina reads are aligned to a reference genome (NC_000913.3), and a list of variants is output that was then analyzed with custom scripting in Python 3.7. Code-base is available online at http://churchlab.github.io/C321evolve/. In brief, variants were culled based on minimum read depth and allele frequencies, multiply-hit genes were identified, cross-contamination events were sorted out, and a final list of genes was assembled that saw at least three unique mutational episodes in any of the four strains. (ii) A custom analytics pipeline was run in parallel to the Millstone analysis, using bowtie2 version 2.1.0 (85) to align reads to the reference genome and Picard version 1.44 to mark duplicate reads. A permissive list of candidate SNPs and indels was generated by applying the GATK UnifiedGenotyper (86) version 2.3 to all samples at once, with the minimum phred-scaled confidence threshold set to its most permissive value, 4.0. The list of candidate mutations was then filtered by excluding all sites with more than one alternate allele and all indels longer than one base pair because these sites are less likely to contain true variants and harder to call confidently. To ensure adequate coverage, it was required that the site: (1) have an average coverage depth of at least 10x across samples, (2) have zero coverage depth in no more than one sample, and (3) have at least ten total reads supporting the alternate allele. Because the samples are haploid, we required the alternate allele to be supported by at least 85% of the reads in at least one sample. Finally, read were filtered on GATK’s quality score and strand bias fields (requiring QUAL > 50 and FS < 40).

### Allele reconstructions

MAGE was used to reconstruct mutations of interest in the ancestral strains. Saturated overnight cultures were diluted 100-fold into 3 mL LB containing appropriate antibiotics and grown at 34 °C until mid-log. The integrated Lambda Red cassette in C321.ΔA derived strains was induced in a shaking water bath (42 °C, 300 rpm, 15 minutes), followed by incubation on ice for at least two minutes. The cells were then made electrocompetent at 4 °C by pelleting 1 mL of culture (16,000x rcf, 20 seconds) and washing twice with 1 mL ice cold deionized water (dH2O). Electrocompetent pellets were resuspended in 50 μL of dH2O containing the desired DNA. For MAGE oligonucleotides, 5 μM of each oligonucleotide was used. Please see Table 2 for a list of all oligonucleotides used in this study. For integration of dsDNA cassettes, 50 ng was used. Allele-specific colony PCR was used to identify desired colonies as previously described (87). Colony PCR was performed using Kapa 2G Fast HotStart ReadyMix following manufacturer protocols and Sanger sequencing was performed by Genewiz to verify strain engineering.

### Doubling time measurement

To assess fitness by measuring doubling times, strains were grown in triplicate in transparent flat-bottom 96-well plates (150 μL, 34 °C, 300 rpm). Kinetic growth (OD_600_) was monitored on Eon plate readers at 5 minute intervals. After blanking reads by subtracting wells containing only media, doubling times were calculated by t_double_ = C*ln(2)/M, where C = 5 minutes per time point and M is the maximum slope of ln(OD_600_). Since some strains achieved lower maximum cell densities, slope was calculated based on the linear regression of ln(OD_600_) through 5 contiguous time points (20 minutes) rather than between two pre-determined OD_600_ values.

## Acknowledgements

We thank Alex LeFell, Seth Shipman, George Chao, Erkin Kuru, Gabriel Filsinger, Gleb Kuznetsov, and Daniel Goodman. This project was graciously funded by the US Dept. of Energy under grant DE-FG02-02ER63445. GMC has related financial interests in ReadCoor, EnEvolv, and GRO Biosciences. For a complete list of GMC’s financial interests, please visit http://arep.med.harvard.edu/gmc/tech.html.

